# *L. pneumophila* resists its self-harming metabolite HGA via secreted factors and collective peroxide scavenging

**DOI:** 10.1101/2023.05.11.540380

**Authors:** Mische Holland, Danielle N. Farinella, Emily Cruz-Lorenzo, Madelyn I. Laubscher, Darian A. Doakes, Maria A. Ramos, Nanami Kubota, Tera C. Levin

**Affiliations:** Department of Biological Sciences, University of Pittsburgh, Pittsburgh, PA; Department of Plant and Microbial Biology, University of California Berkeley, Berkeley, CA; Department of Microbiology and Molecular Genetics, University of Pittsburgh School of Medicine, Pittsburgh, PA

## Abstract

Many pathogenic bacteria, including *Legionella pneumophila*, infect humans from environmental reservoirs. To survive in these reservoirs, bacteria must withstand microbe-on-microbe competition. We previously discovered that *L. pneumophila* can compete with neighboring bacteria via an antimicrobial metabolite called homogentisic acid (HGA) (Levin, Goldspiel, and Malik 2019). Curiously, *L. pneumophila* strains that secrete HGA are not wholly immune to its effects: low-density bacteria are strongly inhibited by HGA whereas high-density cells are tolerant. How do these bacteria tolerate HGA and avoid self-harm during interbacterial competition? Here, we find that HGA toxicity occurs via the production of toxic hydroperoxides and multiple factors facilitate high-density tolerance. First, HGA only becomes fully toxic after >1 hour of oxidation. While this manifests as a delay in killing within well-mixed liquid cultures, in a biofilm environment, this could provide time for HGA to diffuse away before becoming toxic. Second, HGA generates quantities of hydroperoxides that can be collectively scavenged by high-density, but not low-density cells. And third, high-density cells produce one or more secreted factors that are transiently protective from HGA. In combination, we propose that the bacteria are able to deploy HGA to generate a pool of reactive oxygen species surrounding their own biofilms, while maintaining non-toxic conditions within them. Overall, these findings help to explain how broadly toxic molecules can be used as inter-bacterial weapons. They also provide insights about why some of our current decontamination methods to control *L. pneumophila* are ineffective, leading to recurrent disease outbreaks.

## Introduction

As bacteria compete for nutrients and space, many species use antimicrobial compounds to antagonize their neighbors (Ghoul and Mitri 2016). While some of these molecules target only specific strains, those that generate reactive oxygen species (ROS) tend to be broadly active, as ROS can damage DNA, proteins, lipids, and other biological molecules (Vatansever et al. 2013). In oxygenated environments, redox-active secreted compounds, such as phenazines, quinones, and related molecules can generate toxic ROS through redox cycling (O’Brien 1991). Yet despite their toxicity, many bacteria secrete these molecules in high quantities, suggesting that redox-active compounds can confer a number of benefits on the bacteria that produce them, such as improved cellular respiration and improved scavenging of limiting nutrients from the environment (Price-Whelan, Dietrich, and Newman 2006; Zheng et al. 2013; McRose and Newman 2021).

*Legionella pneumophila* is an aquatic opportunistic human pathogen that can cause large disease outbreaks if it colonizes building plumbing systems (Cunha, Burillo, and Bouza 2016). We previously described the secreted quinone HGA (homogentisic acid) as an antimicrobial metabolite used by *Legionella pneumophila* (Levin, Goldspiel, and Malik 2019). In stationary phase, this bacterium secretes abundant (50-500µM) HGA, which can spread across an agar plate to kill neighboring *Legionella* species. We demonstrated that HGA’s oxidation state was relevant for its activity, that reducing agents made HGA inhibition less potent, and that HGA was non-toxic in anaerobic conditions. We therefore proposed that reactive intermediates were inhibitory, although HGA’s mechanism of action was unknown. In addition, we previously observed that *L. pneumophila* could itself be antagonized by HGA and its susceptibility depended on cell density; dense cells (∼2E9 CFU/mL) were insensitive to 24h of HGA treatment but the same strain was extremely sensitive when diluted (∼2E7 CFU/mL) (Levin, Goldspiel, and Malik 2019). HGA susceptibility was particularly strong when cells were exposed to HGA in PBS, which more closely approximates the low-nutrient, aquatic conditions where *Legionella* are found than rich media conditions. In summary, we found that *L. pnuemophila* strains that antagonize neighbors with secreted HGA can themselves be extremely sensitive to this same molecule. How, then, is *L. pneumophila* able to use HGA to kill other bacteria while avoiding self-harm?

Here, we investigate HGA’s mechanism of action and its density-dependent susceptibility. While HGA itself is non-toxic, our findings suggest that it spontaneously generates toxic hydroperoxides over time in aerobic conditions. While dilute *L. pneumophila* are highly sensitive, more dense bacteria can tolerate >1000-fold higher HGA concentrations. Moreover, the threshold separating high-density tolerant cells and low-density susceptible cells is incredibly sharp, separated by only a 2-3x difference in cell density. How is this switch-like difference in susceptibility achieved? First, we find that high-density cells secrete one or more factors (likely small molecule antioxidants) that reduce the amounts of toxic ROS generated by HGA; the secreted factor(s) can transiently protect low-density cells from HGA. Second, HGA generates concentrations of H2O2 that high-density cells can cooperatively detoxify but low-density cells cannot. And third, because HGA takes >1hr to begin to produce toxic levels of ROS in aerobic environments, we propose that this time delay acts as a metaphorical long fuse, allowing HGA to diffuse away from high-density cells in a biofilm before generating toxic ROS. Together, these dynamics permit high-density *L. pneumophila* to secrete abundant HGA without suffering any loss in viability.

## Results

### HGA induces rapid, density-dependent cell death

To precisely define when and how HGA inhibition occurs, we first investigated which cell densities were susceptible to HGA. When we exposed cells across a range of cell densities to 125 µM HGA for 24h, we found that the transition between complete HGA susceptibility and full tolerance was incredibly sharp, spanning only a 2.5- to 3-fold difference in starting cell density (1.35E8 vs. 3.45E8 CFU/mL, Fig 1A). Above the density threshold, *Legionella* were essentially unaffected by HGA, as the number of CFUs at t=0 matched the number we recovered at t=24h (y=x line, Fig 1A). Below the threshold, we did not recover any viable CFUs. We interpret this switch-like behavior to indicate HGA sensitivity is a biologically regulated process, rather than a generic outcome resulting from more cells being present in high-density cultures.

**Figure 1.**
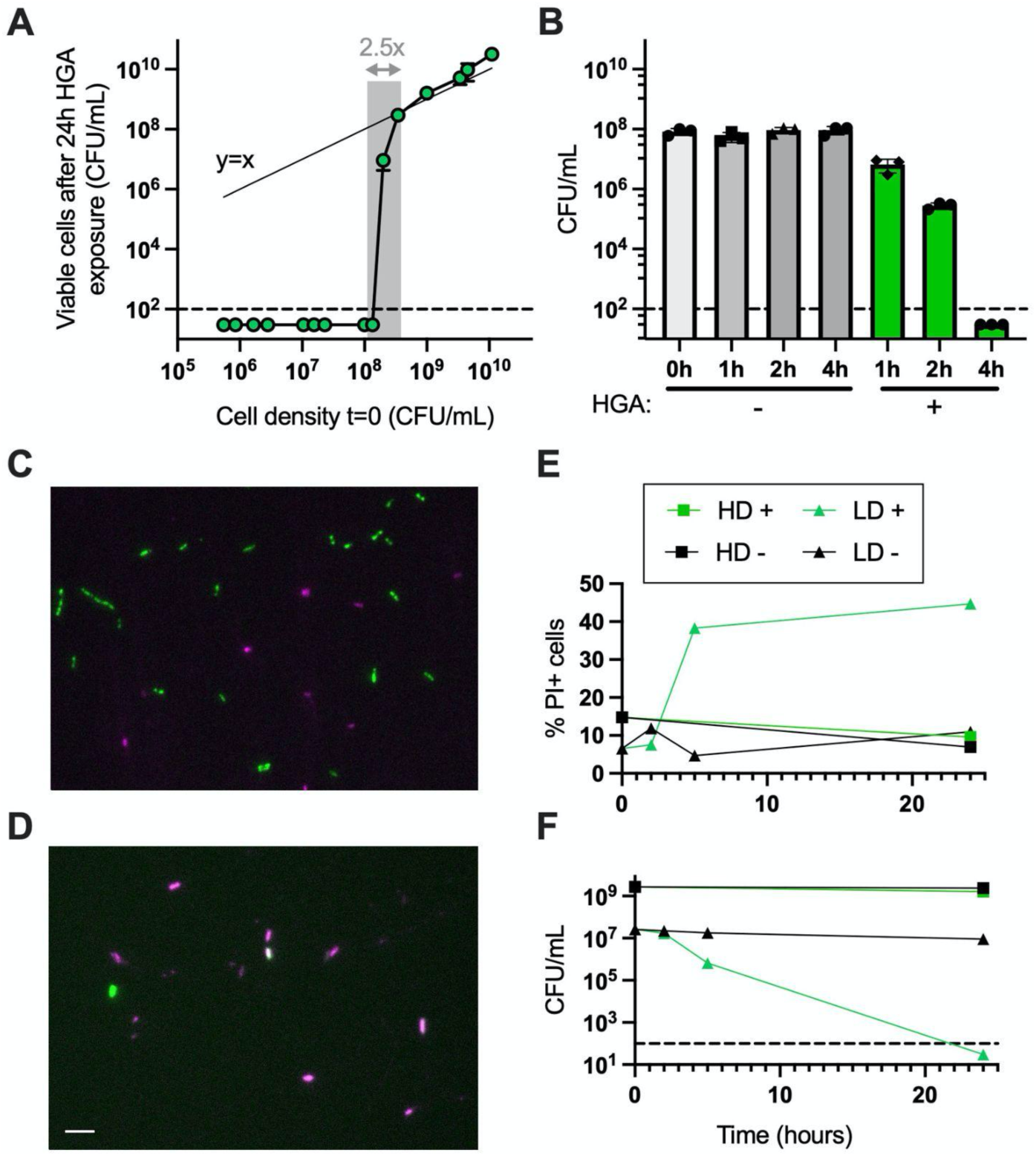
HGA induces rapid, density-dependent cell death. **A)** At high densities, *L. pneumophila* cells are insensitive to HGA; CFUs recovered before and after HGA exposure are similar, matching the y=x line (thin line). However, within only a 2.5-fold change in starting cell density (shaded region), *Legionella* transition from fully tolerant of HGA to completely susceptible. **B)** Low density *Legionella* die rapidly from 125µM HGA, with CFUs unrecoverable after 4 hrs. **C-D)** Representative 24h images of GFP+ *Legionella* exposed to HGA at high **(C)** or low density **(D)** and stained with PI (magenta) to visualize dead cells. Scale bar = 5µm. E) Quantification of PI+ cells and F) viable CFU/mL during microscopy experiment of high- and low-density cells (HD/LD) incubated both with (+) and without HGA (-). Dashed line shows the limit of detection in all experiments.

We also found that the loss of low-density (<7E7 CFU/mL) cell viability occurred quickly. CFUs began to decline after 1h and full inhibition occurred as early as 4h (Fig 1B). The kinetics of inhibition varied depending on culture volume and shaking speed (Sup Fig 1), likely due to the efficiency of oxygenation. To determine if HGA inhibition is bactericidal or if it is driving cells into a viable but non-culturable state, we exposed GFP-expressing *Legionella* to HGA and stained dead cells with propidium iodide (PI) (Fig. 1C-D). We found up to 45% of low-density cells exposed to HGA became PI+. Additionally, the proportion of PI+ cells increased at the same time that these cells experienced a drop in CFUs (Fig. E-F), indicating that the low-density *Legionella* are dying after exposure to HGA. Notably, our % PI+ measurements in this assay are likely an underestimate of total dead cells, as we expect to miss cells that have already lysed. Overall, we conclude that HGA is bactericidal and kills low-density cells rapidly. High-density cells remain unaffected and there is a tight density threshold separating the two outcomes of dramatic cell death vs. complete HGA tolerance.

### Density-dependent susceptibility to HGA and H2O2

Many antibiotics are known to produce an “inoculum effect”, wherein higher antibiotic concentrations are required to kill dense bacteria as compared to dilute bacteria (Soriano, Edwards, and Greenwood 1992; Udekwu et al. 2009). To determine if HGA’s density dependence behaved similarly to inoculum effects, we measured the minimum bactericidal concentration (MBC) for high- and low-density *Legionella* exposed to a variety of antimicrobial compounds in minimal PBS media. Strikingly, over 1,000-fold more HGA was required to fully kill high-density cells (20 mM HGA) than low-density cells (20 µM HGA, Fig. 2A). We observed a similar density-dependence in *Legionella*’s sensitivity to H2O2 (Fig. 2B, 30µM vs. 30mM to kill >99.9% of low- vs. high-density cells).

**Figure 2.**
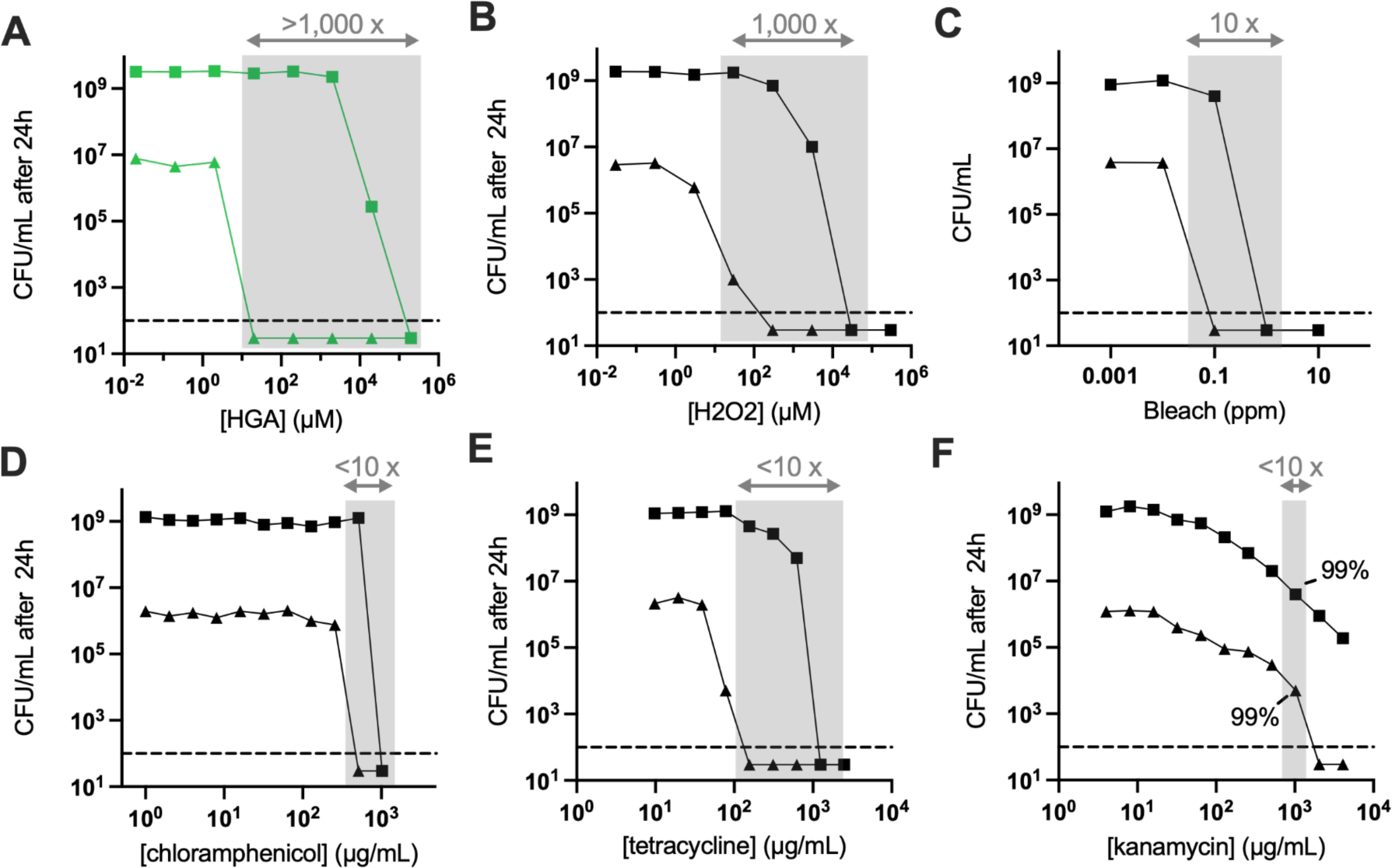
*L. pneumophila* cell density has a large impact on HGA and H2O2 susceptibility. **A**) HGA kills low density bacteria (triangles) at ∼5,000X lower concentration than is required to kill high density cells (squares) after 24h. Shaded regions show the differences in inhibitory concentrations between high and low density *Legionella* for each compound. **B-C**) High density-dependence of *Legionella* susceptibility to H2O2 (**B**) and, to a lesser extent, bleach (**C**). **D-F**) *Legionella* susceptibility to various antibiotics shows a more typical inoculum effect. In F where solubility issues limited our ability to kill high density cells below our limit of detection, we highlight the amount of drug required to kill 99% of the starting bacterial population. All experiments incubated bacteria with antimicrobial compounds in PBS for 24 hours.

In contrast to this density-linked sensitivity, we observed that *L. pneumophila* treated with bleach, multiple antibiotics, and the toxic lipid 4-HNE, exhibited at most a 10-fold difference in the MBCs between high- and low-density *L. pneumophila* (Fig. 2C-F, Sup. Fig 2A). As an additional oxidative stress, we tested *Legionella’s* susceptibility to paraquat, a chemical that generates superoxide. Although previous research found *L. pneumophila* was sensitive to 0.25-0.5mM paraquat in rich media (Sadosky et al. 1994; LeBlanc, Davidson, and Hoffman 2006), in the minimal media used here we found *L. pneumophila* tolerated up to 21mM paraquat with minimal cell death at both high and low cell densities (Sup. Fig 2). We note that the MBCs measured here were higher than previously reported MBCs and MICs for *Legionella pneumophila (Moffie and Mouton 1988; Gibson and Fitzgeorge 1983; Isenman et al. 2018)*. This discrepancy likely results from the fact that antimicrobials are typically added during bacterial log phase growth in chemically complex, rich media, while our treatments are added to cells in minimal media that are no longer growing. For HGA, we previously showed that this compound is <u>more</u> toxic in minimal vs. rich media and this difference is due to high concentrations of cysteine in rich media, which acts as an antioxidant to protect from HGA (Levin, Goldspiel, and Malik 2019). We further found that *Legionella* growth phase had no impact on low-density susceptibility nor high-density HGA tolerance. Overall, *L. pneumophila’s* density-dependent susceptibility to both HGA and H2O2 is several orders of magnitude stronger than the inoculum effect we observed for other antimicrobial compounds (Fig. 2A-B vs 2C-F). When we tested several other species of bacteria (*Legionella micdadei*, *Pseudomonas fluorescens, Klebsiella aerogenes,* and *Bacillus subtilis*) in the same assay, *L. micdadei* and *B. subtilis* were both susceptible to HGA. However, only *L. pneumophila* exhibited a strong density-dependence to both 125µM HGA and 300µM H2O2 (Sup. Fig 2).

### HGA tolerance is independent of the Lqs quorum sensing and HGA synthesis pathways

At high density, bacteria often experience nutrient limitation, slowed growth, and other stresses that could alter bacteria sensitivity to antimicrobials. However, we previously found that *L. pneumophila*’s high-density tolerance of HGA was unaffected by log vs. stationary growth phase, nor was it impacted by the stringent response pathway, which coordinates many stress responses in high-density, stationary phase bacteria (Levin, Goldspiel, and Malik 2019; Molofsky and Swanson 2004). Instead, HGA susceptibility was linked specifically to cell density. Therefore, we next tested whether *Legionella* regulates its density-dependent responses to HGA via candidate quorum sensing pathways. During quorum sensing, bacteria sense their own cell density through the secretion of autoinducer molecules, which accumulate to high concentrations when cells are dense and activate downstream signaling pathways (Abisado et al. 2018; Mukherjee and Bassler 2019). Within the sole characterized *<u>L</u>egionella* <u>q</u>uorum <u>s</u>ensing (Lqs) pathway in *L. pneumophila*, the protein LqsA synthesizes a secreted autoinducer, which is sensed by transmembrane kinases, ultimately altering the activity of the response regulator LqsR (Hochstrasser and Hilbi 2017). We previously found that wild type cells and a Δ*lqsR* mutant had similar HGA susceptibility (Levin, Goldspiel, and Malik 2019), suggesting that this protein is not involved in regulating HGA susceptibility. However, a second response regulator, LvbR, was recently discovered in the Lqs pathway, with some downstream target genes that are not regulated by LqsR (Hochstrasser et al. 2019). Signaling by LvbR, but not LqsR, was also found to be important for *Legionella* development into antibiotic tolerant persister cells (Personnic, Striednig, and Hilbi 2021). To test if LvbR or another part of the Lqs pathway regulates HGA tolerance, we constructed deletion mutants of *lvbR* and *lqsA.* We found that all Lqs mutants had similar HGA susceptibility to wild type (Sup. Fig 3), supporting and strengthening our prior conclusions that density dependent HGA susceptibility is regulated by an independent mechanism.

One secreted molecule that accumulates specifically in high-density *Legionella* cultures is HGA itself. We next considered the possibility that HGA could function as an autoinducer, as secreted quinones have been shown to act as signaling molecules in other bacterial species (Ji et al. 2013). In this scenario, high density cells could begin to secrete HGA, sense the initial, relatively low concentrations of extracellular HGA around them, and activate their protective downstream pathways before releasing subsequent, toxic levels of HGA. If this hypothesis is correct, we predicted that *Legionella* mutants that are unable to secrete HGA would be susceptible to exogenous HGA, even at high cell density. Contrary to this expectation, we observed that a mutant that secretes no HGA (Tn:*hisC2*, (Levin, Goldspiel, and Malik 2019)) showed similar patterns of susceptibility to exogenous HGA as both wild type cells and *ΔhmgA* mutants that secrete excess HGA (Sup. Fig 3). We therefore concluded that *Legionella*’s density-dependent susceptibility is unrelated to both HGA biosynthesis and to Lqs pathway signaling.

### *L. pneumophila* RNA-seq reflects low-density cell oxidative stress from HGA

To identify genes that could underlie *Legionella*’s density-dependent response to HGA, we performed RNA-seq on high- and low-density cells, both with and without HGA exposure (i.e. HD+, HD-, LD+, and LD-conditions). Because LD+ cells begin to experience substantial cell death after 1-2h of HGA exposure (Fig. 1B), we analyzed RNA from each condition at the t=1h time point. By principal components analysis, the transcriptional profiles from HD samples were clearly separated from LD samples along PC1, while LD- and LD+ samples differed along PC2 (Sup. Fig 4). The LD+ samples also had the most within-condition variation. Because LD+ cells experience a steep decline in CFUs between t=0-2h, we expect that the differences among replicates reflect slight experimental variations in timing or degree of ROS generation from HGA. In agreement with the PCA, correlation matrices of the normalized read counts within conditions showed low variation between replicates within each condition except for the LD+ condition (Sup. Fig 4).

From the RNA-seq data, we first asked whether high-density cells produced an inducible, protective response to HGA by comparing HD- and HD+ samples. Contrary to this hypothesis, we found that <u>zero</u> genes were differentially expressed when high-density cells were exposed to 125µM HGA (Sup. Table 1). Thus, in addition to high-density cells’ HGA tolerance (i.e. they have no decline in CFUs, Fig 1B), these cells do not appear to activate any stress or signaling responses to HGA at the transcriptional level.

In contrast, low-density cells robustly upregulated 8 genes when exposed to HGA, most related to oxidative stress (Fig. 3A). These included *sodC* (*lpg2348*), a copper-zinc superoxide dismutase, and *ahpC2D* (*lpg2349 & 2350*), which encode an alkyl hydroperoxidase and reductase that together scavenge hydroperoxides such as H2O2 (LeBlanc, Davidson, and Hoffman 2006). Also upregulated was an operon containing *pirin* and *acpD* orthologs (*lpg2286 & 2287*). While both genes are uncharacterized in *Legionella*, pirins are found across bacteria and eukaryotes where they play diverse roles, including acting as redox-sensing transcriptional regulators (Liu et al. 2013) or directly degrading aromatic antimicrobial compounds (Adams and Jia 2005). AcpD is an azoreductase, acting on aromatic nitrogen-containing compounds (Nakanishi et al. 2001). The final three upregulated genes were related to iron acquisition. The *feoAB* operon (*lpg2658/2657*) mediates the transport and uptake of Fe^2+^, while *frgA (lpg2800*) synthesizes a siderophore for uptake of Fe^3+^ (Cianciotto 2015). Oxidative stress has previously been shown to induce an iron starvation response via oxidation of Fe^2+^ and subsequent release of the Fur repressor (Varghese et al. 2007), consistent with the gene expression observed here. Notably, the 8 genes upregulated in LD+ conditions represent only a subset of *Legionella’s* oxidative stress response genes. In none of the RNA-seq conditions did we observe differential expression of *Legionella*’s other superoxide dismutase *sodB*, nor the other alkyl hydroperoxidase *ahpC1*, nor any previously characterized catalase or *oxyR* regulator genes (Fig. 3C, Sup. Table 1). Similarly, none of the *Legionella* quorum sensing (*lqs*) pathway genes nor the HGA biosynthesis genes were differentially expressed (Sup. Fig 4).

**Figure 3.**
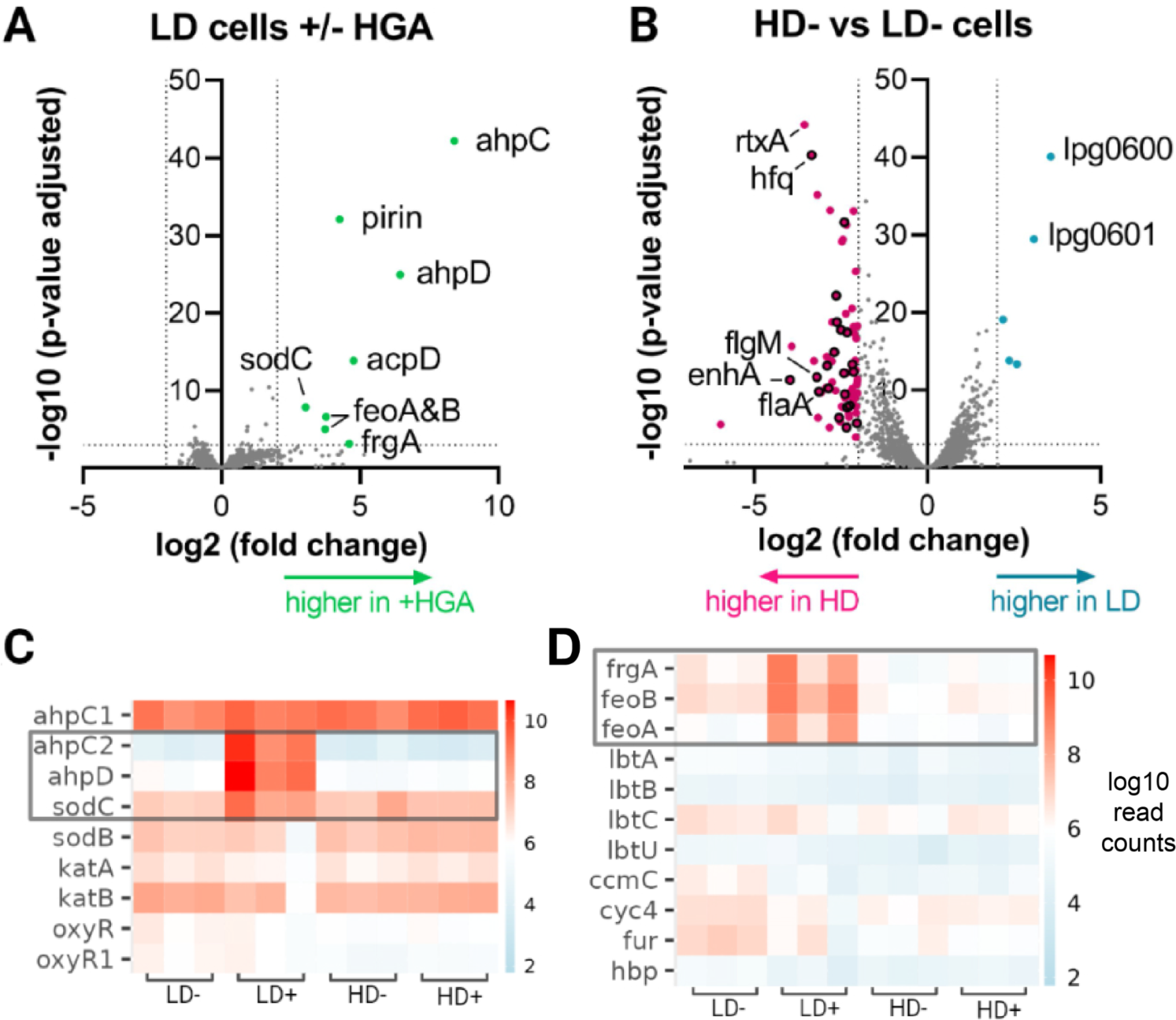
Differentially expressed genes identified by RNA-seq **A)** Volcano plot showing differential gene expression in low-density *L. pneumophila* with and without HGA exposure. Dotted lines indicate significance thresholds and significantly different genes are colored. **B)** Differential expression between high-density and low-density cells in the absence of HGA. Genes related to flagellar function are outlined in black. **C-D)** Heatmaps of RNA expression level for key genes in the oxidative stress **(C)** and iron scavenging pathways **(D)**. Grey box indicates genes that were significantly upregulated in LD+ cells.

Because high-density *Legionella* show no change in viability or gene expression when exposed to HGA, we reasoned that the mechanisms of high-density tolerance are likely due to differences between high- and low-density cells prior to HGA exposure. Between HD- and LD-conditions, there were 78 genes significantly upregulated in high-density cells and 5 genes upregulated in low-density cells. *Legionella* exhibits a biphasic lifestyle and is known to upregulate flagellar biosynthesis and virulence gene expression in stationary phase, downstream of the stringent response pathway (Albert-Weissenberger et al. 2010; Appelt and Heuner 2017). The predominant transcriptional profile of genes upregulated in HD-cells was consistent with this flagellar regulon activation. Twenty-one of the 78 upregulated HD-genes were flagellar (in flagellar gene classes II, III, and IV) or previously characterized as part of the *fliA* flagellar regulon (Albert-Weissenberger et al. 2010). If we relax our significance threshold to p<0.05, 60% of all flagellar-related genes in the genome (34 of 57) were upregulated in HD-as compared to LD-conditions (Sup. Table 1). Relatedly, 9 characterized virulence genes (i.e. toxins or effectors) were upregulated in HD-conditions. While this was the major transcriptional signature, we previously showed that HGA susceptibility was unaffected by mutations to the stringent response pathway (Levin, Goldspiel, and Malik 2019), which are expected to perturb activation of the flagellar regulon (Appelt and Heuner 2017). Therefore, we predict that HGA tolerance is more likely to be conferred by one or more of the remaining, non-flagellar genes upregulated in high-density cells. Of these, there was no obvious pattern, partly due to the fact 47% of the remaining genes were hypothetical and uncharacterized, mostly small proteins that are predicted to be unstructured by AlphaFold. There were also two heat shock proteins belonging to the HSP20 family (*lpg2191 & 2493*) as well as several predicted transcriptional regulators that could mediate the expression of genes at high density (e.g. *lpg0476, 0586, 2008, 2181,* and *2145*).

Within low-density cells, the two most upregulated were *lpg0600,* an Rrf2 family transcription factor, and *lpg0601*, an ABC transporter. Notably, these genes lie closely upstream of *lpg0607*, a gene that was repeatedly disrupted in a screen for HGA-tolerant mutants in our previous study (Levin, Goldspiel, and Malik 2019), although *lpg0607* itself was only weakly upregulated in low-density cells (1.8-fold higher, p=0.043). It is possible that regulation of this operon modulates HGA sensitivity in low-density bacteria. Overall, the RNA-seq results revealed many genes differentially regulated between high- and low-density cells, as well as specific oxidative stress responses in HGA-exposed, low-density *Legionella*.

### HGA gradually produces hydroperoxides, which can be quenched by high-density cells

Because H2O2 and HGA displayed a strikingly similar density-dependence in *L. pneumophila* (Fig. 2A-B) and because *ahpC2D*, genes used to detoxify hydroperoxides, were highly upregulated in LD+ cells (Fig. 3A), we next hypothesized that HGA kills low-density *L. pneumophila* via generation of H2O2 or a related hydroperoxide. To test this hypothesis, we used the reagent Amplex Red, which generates the fluorescent compound resorufin in the presence of hydroperoxides (Zhou et al. 1997). Previously, we found that HGA was non-toxic in anaerobic conditions (Levin, Goldspiel, and Malik 2019), suggesting that toxic compounds are produced upon oxidation. Consistent with this finding and with the timing of HGA killing of low-density cells (Fig. 1B), we saw that synthetic HGA incubated in shaken test tubes gradually produced abundant fluorescent signal. When compared to a standard curve of H2O2, 125µM HGA generated an equivalent signal of 130µM H2O2, reaching hydroperoxide levels that are toxic to low-density cells after 1hr (Fig 2B & 4A). As we found that high-density cells could tolerate H2O2 but low-density cells died between 30-3000 µM H2O2 (Fig. 2B), the fact that HGA generates hydroperoxides within this critical range may contribute to *L. pneumophila*’s density-dependent susceptibility to HGA.

However, it is also possible that HGA generates different types or concentrations of reactive oxygen species in the presence of bacteria. To test this hypothesis, we incubated HGA alone or in combination with *L. pneumophila* at high- or low-density for 4hrs and measured hydroperoxide concentrations with Amplex Red. We saw that HGA alone generated the equivalent of 92 µM H2O2 and only a slightly lower amount (52 µM) in the presence of low-density cells. However, when HGA was incubated with high-density *L. pneumophila*, the Amplex Red signal was dramatically reduced (0.8µM, Fig. 4B). This behavior was not unique to HGA, as high-density cells also depleted H2O2 during co-incubation (Fig. 4B).

**Figure 4.**
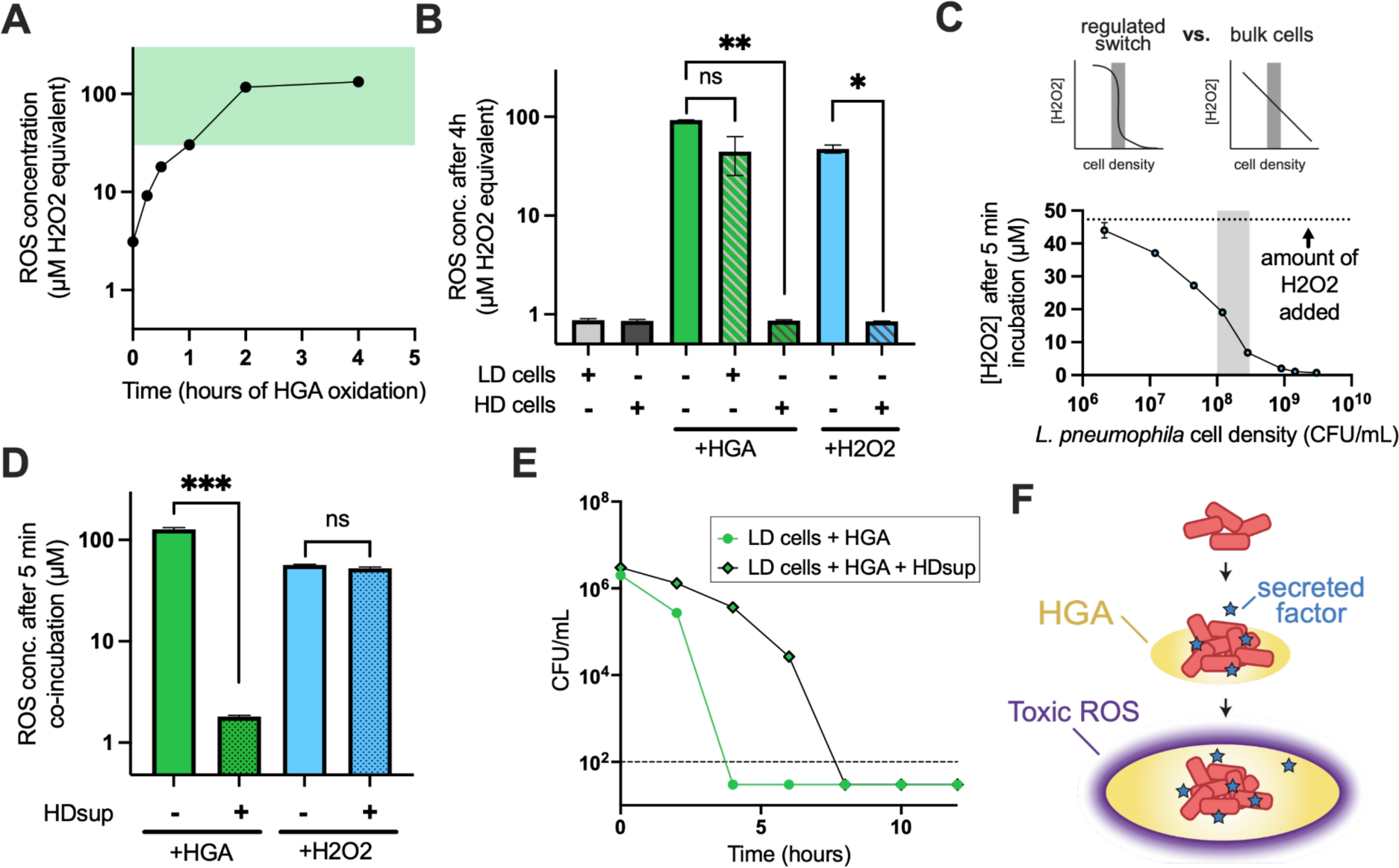
Dynamics of HGA toxicity, collective ROS detoxification, and secreted anti-ROS factors allow high-density cells to survive HGA exposure. **A)** Generation of ROS upon oxidation of 125µM HGA, measured with Amplex Red. Green shaded region marks H2O2 concentrations that were toxic to low-density, but not high-density cells in Figure 2. **B)** High-density (HD) L. pneumophila reduce the amount of ROS present in HGA- or H2O2-treated cultures, but low-density (LD) cells do not. * = p<0.05 and ** = p<0.005 by Welch’s t-test. **C)** To test if the reduction in H2O2 in the presence of HD cells in panel B was due to the bulk action of many cells vs. a density-dependent switch as we saw for HGA, L. pneumophila of many cell densities were incubated in the presence of 50µM H2O2 for 5min, then the H2O2 remaining was quantified with Amplex Red. The reduction in H2O2 was correlated with cell density but did not exhibit regulated switch behavior. Shaded region shows the cell density threshold observed for HGA in Figure 1A. **D)** High-density supernatants (HDsup) can reduce the Amplex Red signal from HGA (p<0.0005) but has little impact on the H2O2 signal. **E)** Incubation of low-density cells with HDsup provides transient protection from HGA. **F)** Proposed model of HGA activity in biofilms. HGA is secreted by high-density cells and only form toxic levels of ROS after a time delay and spread of HGA. Toxic ROS levels near the high-density cells are kept low via both cooperative ROS detoxification and secreted factors that transiently quench ROS production from HGA.

H2O2 is cell-permeable, and it has previously been shown that the combined actions of bacteria scavenging intracellular H2O2 can collectively reduce local H2O2 concentrations (Ma and Eaton 1992; Choudhary et al. 2023). Because cells at higher density should be better at scavenging H2O2, we next investigated whether this phenomenon was sufficient to explain the density-dependent susceptibility to HGA. If so, we predicted that there would be a switch-like transition between 1E8 to 3E8 CFU/mL (the density threshold defined in Fig. 1A), where cultures above this threshold would scavenge H2O2 much more efficiently than bacteria below the threshold (see ‘regulated switch’, Fig. 4C). Alternatively, if high-density H2O2 scavenging is simply related to the activity of additional cells, we predicted that cells at higher density would scavenge H2O2 without any apparent density threshold (see ‘bulk cells’, Fig 4C). When we briefly incubated cells at different densities with 50µM H2O2, we observed that higher cell densities reduced the concentration of H2O2 more readily, but this occurred without any apparent density threshold (Fig. 4C). Therefore, while high-density cells exhibit enhanced H2O2 scavenging as expected, this phenomenon is not sufficient to explain HGA’s density-dependent susceptibility.

### High-density cells secrete a transiently protective molecule

We next asked whether the high-density cells’ quenching activity could be enhanced by a secreted factor. We generated secreted supernatants from high-density cells (HDsup) by incubating dense *L. pneumophila* in PBS for 1 hour and then centrifuging and filter sterilizing the sample through a 0.2µm filter to remove bacteria. To generate HGA-derived ROS, we oxidized HGA alone for 4 hours. When we then incubated the high-density supernatants with HGA, we found that the HDsup could largely quench the Amplex Red signal (Fig. 4D). Unlike the activity from live cells, HDsup incubated with H2O2 did not alter the H2O2 measurement. This suggests that high-density cells rapidly secrete one or more factors that are protective from HGA. Because HDsup does not substantially alter H2O2 readings, we infer that the decline in fluorescence is due to ROS quenching and that HDsup does not interfere with the Amplex Red assay. Moreover, because HDsup does not detoxify H2O2 but it does alter ROS generated from HGA, we conclude that HGA produces one or more toxic ROS that reacts with the Amplex Red reagent but is distinct from H2O2.

If high-density cells resist HGA via a secreted factor, we predicted that HDsup would protect low-density cells from HGA. Indeed, we found that low-density cells incubated with HDsup experienced delayed cell death from HGA, extending the time to death by at least 4 hours (Fig. 4D). This effect was transient, and the cells still died by t=8 hours. Nevertheless, these experiments reveal that HGA generates quantities of ROS that are toxic to low-density cells, but that high-density cells can transiently quench or scavenge extracellular HGA-derived ROS via a secreted, protective factor.

## Discussion

We previously discovered that *L. pneumophila* can use HGA as a secreted, antimicrobial metabolite to antagonize neighboring *Legionella* bacteria (Levin, Goldspiel, and Malik 2019). Here, we have shown that HGA is bactericidal, clarified its mechanism of action, and uncovered mechanisms that mediate *L. pneumophila*’s unusual, density-dependent susceptibility to this molecule. We find that low-density *L. pneumophila* cells are over 1,000x more sensitive to HGA than high-density bacteria (Fig. 2). This sensitivity is mirrored in *L. pneumophila*’s responses to H2O2 but is distinct from other antimicrobials or oxidative stressors (Fig. 2, Sup Fig 2). Based on these observations, as well as the upregulation of alkyl hydroperoxidases in LD+ samples (Fig. 3A), and the measurement of HGA-derived ROS using Amplex Red (Fig 4C), we propose that HGA generates toxic hydroperoxides (distinct from H2O2), which kill low-density *L. pneumophila*. Supporting this conclusion, we found that *L. micdadei* and *B. subtilis* are sensitive to HGA but relatively insensitive to H2O2 (Sup Fig. 2). Further ruling out H2O2 as an intermediate, *L. micdadei tatlock* is also a naturally catalase-positive strain (Pine et al. 1984), yet it remains susceptible to HGA and we found previously that catalase was not protective against HGA (Levin, Goldspiel, and Malik 2019). While Amplex Red is specific for hydrogen peroxide over superoxide, other compounds including peroxynitrite (Dębski et al. 2016) and some organic hydroperoxides (Lietz 1958) have previously been found to react in the Amplex Red/horseradish peroxidase assay, which could mediate HGA toxicity.

We also found that HGA susceptibility in *L. pneumophila* follows an extremely sharp density threshold; the only difference between bacteria that are fully impervious to HGA vs. those that are completely susceptible is a 2-to-3-fold difference in starting cell density (Fig. 1). We interpret this sharp transition to be the result of active, density-dependent regulation. So, what explains this switch-like difference in HGA susceptibility in *L. pneumophila?* Our findings suggest that this phenomenon is distinct from the ‘inoculum effect’ typically seen for antibiotics (Fig. 2), the HGA biosynthesis pathway, and the sole, known quorum sensing pathway of *Legionella* (Lqs) (Sup Fig. 3).

Instead, our results suggest that at least three phenomena contribute to high-density HGA tolerance. First, and best characterized previously, the combined actions of bacteria scavenging toxic hydroperoxides within their own cells can collectively reduce environmental peroxide concentrations and this process occurs more quickly when more bacterial cells are present(Ma and Eaton 1992; Choudhary et al. 2023). We find that HGA produces hydroperoxide concentrations that can be quickly, collectively scavenged by high-density but not low-density *L. pneumophila* populations (Fig. 4A-C). However, this collective scavenging alone is not sufficient to explain HGA’s density-dependent susceptibility.

Second, high-density *L. pneumophila* secrete unknown factors that partially quench HGA-derived ROS (Fig. 4D) and provide transient protection from HGA (Fig. 4E). This transient effect was not detectable in our prior study (Levin, Goldspiel, and Malik 2019), because those experiments only tested samples after 24 hours of HGA treatment. We hypothesize that the transient protection arises because the protective factor is degraded, used up, or oxidized as it interacts with ROS. However, if high-density cells continue to secrete fresh protective factors, these molecules could provide longer-lasting protection.

Third, we propose that the time delay between HGA secretion and the production of toxic ROS could provide an additional level of protection in spatially structured environments. We initially discovered that HGA could kill bacteria in an agar plate assay wherein HGA produced by a *L. pneumophila* biofilm spread across the agar and killed neighboring bacteria (Levin, Goldspiel, and Malik 2019). Because we found HGA is non-toxic when anaerobic (Levin, Goldspiel, and Malik 2019), the HGA would likely remain non-toxic at the center of the producing biofilm, only beginning to oxidize and produce ROS once it reached the biofilm periphery. Even in highly oxygenated environments, we found that it takes >1hr for HGA to generate toxic levels of hydroperoxides (Fig. 4A), during which time HGA would continue to diffuse away from the high-density, HGA-producing cells. This delay in HGA toxicity, combined with local ROS scavenging and quenching near high-density cells could create a low ROS zone near the site of secretion and abundant ROS at more of a distance. In this “toxic moat” model, ROS would surround, but not directly impact, high-density HGA-producing cells (Fig. 4F).

What is the protective factor that is secreted from high-density cells? Typically, bacteria counteract high concentrations of hydroperoxides with catalase enzymes and lower concentrations with peroxidases (Mishra and Imlay 2012). We observed that low-density cells upregulated *ahpC2D* peroxidases upon HGA exposure (Fig. 3A). However, as neither peroxidases, catalases, nor any other typical ROS detoxifying protein was differentially regulated in HD- vs. LD-cells (Fig. 3C), we expect these enzymes are unlikely to be responsible for the protective activity in high-density supernatants. Other secreted factors such as antioxidants could be responsible instead. Indeed, as we previously saw that reducing agents such as cysteine, glutathione, and DTT were protective from HGA (Levin, Goldspiel, and Malik 2019), similar molecules could provide the transient protective activity we observe in high-density supernatants.

Finally, these findings have potential relevance for public health. Because *L. pneumophila* within the built environment can cause serious disease outbreaks, methods of disinfection are of primary interest (Carlson et al. 2020; Casini et al. 2017). Many of the most common methods used, such as chlorine, chloramine, hydrogen peroxide, and ozone, act through the induction of oxidative damage. Our results suggest that *L. pneumophila* susceptibility to certain oxidative stressors strongly depends on cell density, particularly in low-nutrient, aquatic environments. Future decontamination approaches should take into account the fact that high-density bacteria (e.g. which may be found in biofilms) may be many orders of magnitude less susceptible to oxidizing agents than the low-density bacteria typically tested in the lab. For example, some decontamination efforts have used hydrogen peroxide well below the 30mM concentration we find is necessary to fully kill high-density cell populations (Casini et al. 2017). High-density *L. pneumophila* that are insensitive to disinfection methods could form a persistent subpopulation that would be poised to recolonize water systems and contribute to the recurrent disease outbreaks caused by this pathogen (Berjeaud et al. 2016).

## Methods

### Strains and mutant construction

As in Levin et al., 2019, our wild type *Legionella pneumophila* (*Lp*) strain was KS79 and mutants were constructed in this genetic background by allelic exchange using pLAW344. Briefly, we cloned these regions into pLAW344 via Gibson assembly, then transformed into *Lp*, selected for plasmid integration on BCYE-chloramphenicol plates, and counter-selected to create the knock out on BCYE-sucrose plates. To create the *lvbR* knockout strain, the 5’ genomic region was cloned using primers TL180_LvbR_P1_oH1 (acggtatcgataagcttga gcgtgctgattggtcc) and TL181_LvbR_P2_oH1 (ggtttatgaggctga ttgttttttcattc), while the 3’ region was cloned using TL182_LvbR_P3_oH2 (gaatgaaaaaacaa tcagcctcataaacc) and TL183_LvbR_P4_oH2 (cgctctagaactagtggatc cgcagtgccagtcatgac). We used an identical protocol to create the *lqsA* mutant, cloning the 5’ region with primers TL186_LqsA_P1_oH1(acggtatcgataagcttga aatcccctgctccccaaaatag) and TL187_LqsA_P2_oH1 (cgccaataattagaagtcgtg ccataactcagattcttttcc) and cloning the 3’ region with primers TL188_LqsA_P3_oH2 (ggaaaagaatctgagttatgg cacgacttctaattattggcg) and TL189_LqsA_P4_oH2 (cgctctagaactagtggatc atgcaacatttctcaatacc). The GFP-positive *Lp* in the PI microscopy assay were generated by transforming KS79 with the pON-GFP constitutive expression plasmid (Gebhardt, Jacobson, and Shuman 2017).

The other bacterial strains and species used were: *L. micdadei* tatlock (Hébert, Steigerwalt, and Brenner 1980; Garrity, Brown, and Vickers 1980), *Pseudomonas fluorescens* SBW25 (Rainey and Bailey 1996), *Klebsiella aerogenes* DBS0305938 (Dicty Stock Center ID), and *Bacillus subtilis* IAI (WT strain 168 *trpC2*) (Zeigler et al. 2008). *L. micdadei* was cultured on BCYE plates and liquid cultures were grown in AYE, the same culture conditions used for *L. pneumophila*. *P. fluorescens* and *B. subtilis* were cultured at 28C and 37C respectively, on LB plates and liquid cultures were grown in LB. *K. aerogenes* was cultured at 37C on SM/5 plates and liquid cultures were grown in SM media.

### HGA and other antimicrobial susceptibility assays

HGA susceptibility assays were performed in PBS as previously described. Briefly, our core assay was performed with washed bacteria resuspended in 3mL PBS at 3E9-1E10 CFU/mL for high density conditions and 1E7-7E7 CFU/mL for low density conditions, +/-125µM HGA. Cells were plated onto BCYE after 0-24h of incubation and colonies on plates were counted after 2-3 days of 37C incubation for CFU viability.

Density dependent susceptibility to various ROS stress and antibiotic drugs were determined using a standard MBC99 protocol (Hacek, Dressel, and Peterson 1999). High and low density cells were mixed with drug in 200uL of 1X PBS and incubated at 37C in a shaker. Each condition was plated immediately for CFUs on BCYE and plated again after 24 hrs of 37C incubation. The MBC99 or minimum amount of drug required to kill ≥99% of the bacteria in the untreated controls was determined using CFU/mL counts from each drug condition. The antibiotics tested were Kanamycin [4-4096 µg/mL, Thermo Fisher #BP906-5], Chloramphenicol [1-1024 µg/mL, Thermo Fisher BP904-100], and tetracycline [9.76-2500 µg/mL, Sigma #87128-25G]. The ROS producers included H2O2 [0.03-300,000 µM, Fisher Scientific #H325], 4HNE [0.1-1000µM, Cayman Chemical #321001], and paraquat [5.2-21 mM, Sigma #856177]. Most compounds were tested in 2-fold dilutions across the concentration range, while HGA and H2O2 were diluted 10-fold. All compounds were tested in shaken 96-well plates except for H2O2.To avoid volatile H2O2 from cross-reacting between conditions, 1mL cultures were exposed to this compound in separate glass test tubes and culture in a roller drum.

### PI staining and quantification

During the HGA susceptibility assay, cell viability was determined by plating for CFUs and imaging propidium iodide-stained bacterial cells at each timepoint. To define cell boundaries, we used a KS79 *Lp* strain carrying the pON-GFP plasmid to constitutively express GFP (see ‘Strains and mutant construction’). Cells at each timepoint were incubated with 2 µg/mL propidium iodide [Thermo fisher, #P1304MP] for 5 min in the dark just before imaging to stain permeable bacterial cells. To image the cells, stained cell samples were spotted onto GeneFrame agar pads [Fisher scientific #AB-0576] and visualized on a Nikon TE2000 inverted scope using a 60x oil immersion lens in brightfield, red (560 nm), and green (488 nm) channels. For each condition and time point, at least 3 fields of view were quantified, or a minimum of 60 cells each. The data presented here comes from one of two independent experimental replicates.

Images were single-blinded and manually counted for quantification. To determine cell count of green and red cells, the collected image was split into separate color channels using Fiji (Schindelin et al. 2012). Cells (defined as 0.1-10 micron^2^ round particles) were identified using the GFP signal. Cells were defined as PI-positive if they appeared in both red and green channels or if they were visible only in the red channel.

### RNA-seq sample preparation and analysis

The samples prepared for RNA-seq were high- and low-density KS79 cells, with and without HGA. RNA was extracted after 1hr HGA exposure using the RNeasy Mini Kit (Qiagen #74104) as directed, with the addition of a bacterial lysis step using 50 uL 10 mM TrisCL, 1 mM EDTA, 1 mg/mL lysozyme prior to the addition of RLT buffer. RNA samples were treated with TURBO DNase (Fisher scientific #AM1907) to remove contaminating DNA. RNA quality was assessed using the Agilent Tapestation for concentration >50 ng/uL and RIN>6 prior to sequencing. Sequencing libraries were prepared using Illumina Stranded RNA library preparation with RiboZero Plus rRNA depletion (Illumina #20040529).

Each library was sequenced with 50 bp paired-end reads, to a depth of ∼12M reads per sample. Raw sequencing reads were trimmed and filtered using trimmomatic (Bolger, Lohse, and Usadel 2014). After trimming for quality, any reads less than 40 bp and average quality less than 30 were dropped. Trimmed filtered reads were mapped to the reference *Lp* genome (Genbank accession GCA_000008485.1) using kallisto (Bray et al. 2016). The mapped reads were analyzed for differential expression using the DESeq2 R package and a p-value cutoff of 0.05 (Love, Huber, and Anders 2014). DESeq2 normalized read counts across samples and performed the default B-H false discovery rate correction (Benjamini and Hochberg 1995) on p-values to produce the adjusted p-values. We analyzed differential expression according to sample density (high vs low) and HGA (present or absent). Differentially expressed genes were defined as genes with >2-fold change in expression and an adjusted p-value <0.05. To validate that experimental conditions correlated well within replicate, normalized counts data was plotted on a scatter matrix using the pairs R function. To assess variability in our data, we used the vst and plotPCA functions of DESeq2. The differential expression of candidate DEGs and pathways of interest were displayed in heatmaps using ggplot2 v3.3.3 (Wickham 2016). *Legionella* genes involved in iron acquisition and HGA production were defined based on (Cianciotto 2015). Genes involved in ROS response were based on (St John and Steinman 1996; Bandyopadhyay and Steinman 1998; LeBlanc, Davidson, and Hoffman 2006; LeBlanc et al. 2008). Genes involved in the Legionella quorum sensing pathway were based on (Personnic, Striednig, and Hilbi 2018; Hochstrasser et al. 2019) and those in the flagellar regulon were based on (Albert-Weissenberger et al. 2010).

### Preparation of low- and high-density *Lp* for Amplex Red assay

For the experiments in Figure 4B and C, we used a *hisC2*:Tn *Lp* strain from (Levin, Goldspiel, and Malik 2019) that does not secrete HGA. For Fig. 4B, high and low density *Lp* bacteria were washed in PBS and mixed with 125µM HGA, 50μM H2O2 solution, or PBS alone as a control. Each condition was incubated for 4 hours at 37C in a roller drum. After 4 hours each condition was sampled for Amplex Red assays (See “ROS measurements” below). In Fig. 4C, *Lp* bacteria were resuspended in PBS to a density of ∼5e9 cfu/mL, then serial, two-fold dilutions were created down to a density of 3.9e7 cfu/mL. Each culture was then treated with H2O2 at a final concentration of 50µM. Finally, experimental tubes were briefly vortexed and sampled for Amplex Red readings. Samples were read in parallel to an H2O2 standard curve to estimate hydroperoxides present in experimental samples.

### Preparation and activity of *L. pneumophila* supernatants

Supernatants from high-density cells (HDsup) were prepared by washing stationary phase *L. pneumophila* in PBS and incubating 1.5mL of 4-9E9 CFU/mL cells in a rolling-drum incubator at 37C for 1 hour. Supernatants were harvested by centrifugation followed by filter sterilization with a 0.22 µm filter. To test if HDsup could quench HGA-derived ROS or H2O2, we first incubated 125µM of HGA or 50µM of H2O2 in PBS at 37°C for 4 hours in a rolling drum incubator to fully generate ROS. We then mixed 150uL of the pre-oxidized HGA with 2.85mL of HDsup or PBS. The 50µM H2O2 samples were compared with 50µM final in HDsup (i.e. 150uL of 1mM H2O2 + 2.85mL HDsup). All conditions were then briefly vortexed, then sampled for Amplex Red readings after 5min co-incubation.

To test if supernatants impacted bacterial susceptibility to HGA, 1.5mL of *L. pneumophila* cells (at high or low density) were added to 1.35mL HDsup and 150ul of HGA (final HGA concentration 125µM).

### ROS measurements via Amplex Red assay

To quantify the hydroperoxides present in conditions treated with HGA, H2O2, or *L. pneumophila* bacteria, we used the reagent Amplex Red (Thermo Fisher #A12222), which generates a fluorescent compound in the presence of hydroperoxides. Amplex red solution consisted of 50uM of Amplex red and 0.1U of Horseradish peroxidase (Sigma #77332), which was incubated for 10 minutes at 37C. Then, 100ul of Amplex reaction solution was added to 20ul of experimental culture in an opaque 96-well plate. Fluorescence was measured after 5 minutes with a Agilent Biotek Cytation 5 plate reader at 590nm using an excitation wavelength of 530nm. Fluorescence values were converted to H2O2 concentrations using a standard curve of known H2O2 concentrations in parallel.

### Accessions/Data Availability

RNA-seq read archive (SRA): PRJNA943215

RNA-seq read processing and analysis scripts: https://github.com/mliannholland/hga_rnaseq

## Acknowledgements

We thank Ed Culbertson, Jim Imlay, and Jason Yang for useful discussions and suggestions. This work was supported by funding from the Charles E. Kaufman foundation and from the National Institutes of Health.

## Supplemental Figures

**Supplemental Figure 1.**
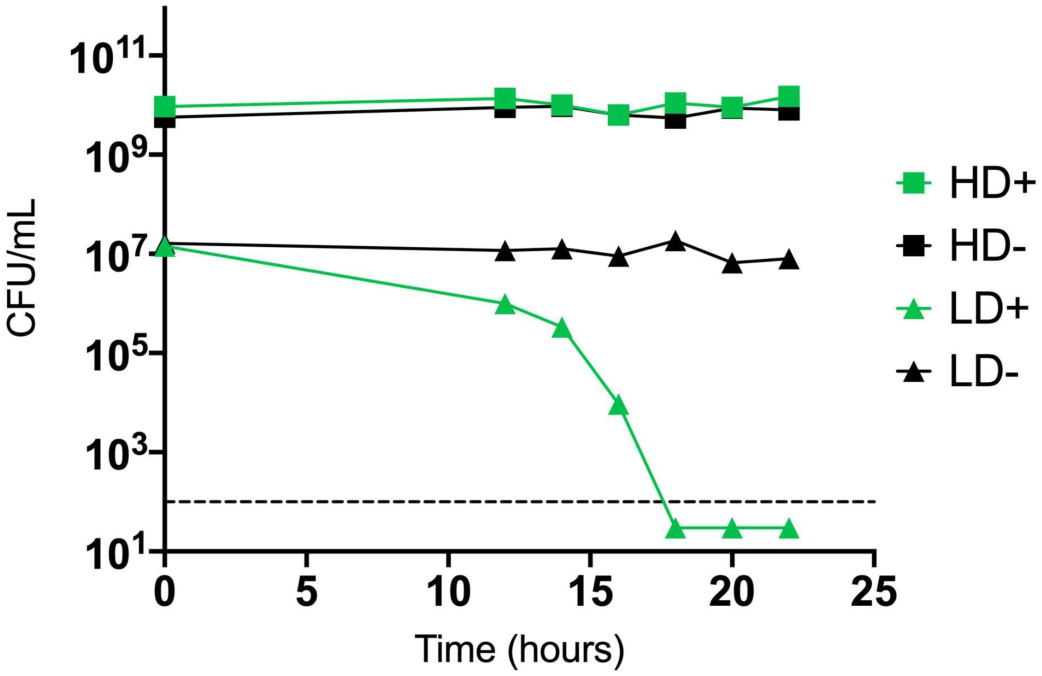
HGA-mediated killing depends on culture volume and shaking conditions. Low-density cells exposed to HGA (LD+ cells) die after 16-18 hours when in 100uL volumes in a 96-well plate, shown here. This is considerably slower than HGA-mediated killing in 3mL culture tubes within a roller drum (Figure 1B).

**Supplemental Figure 2.**
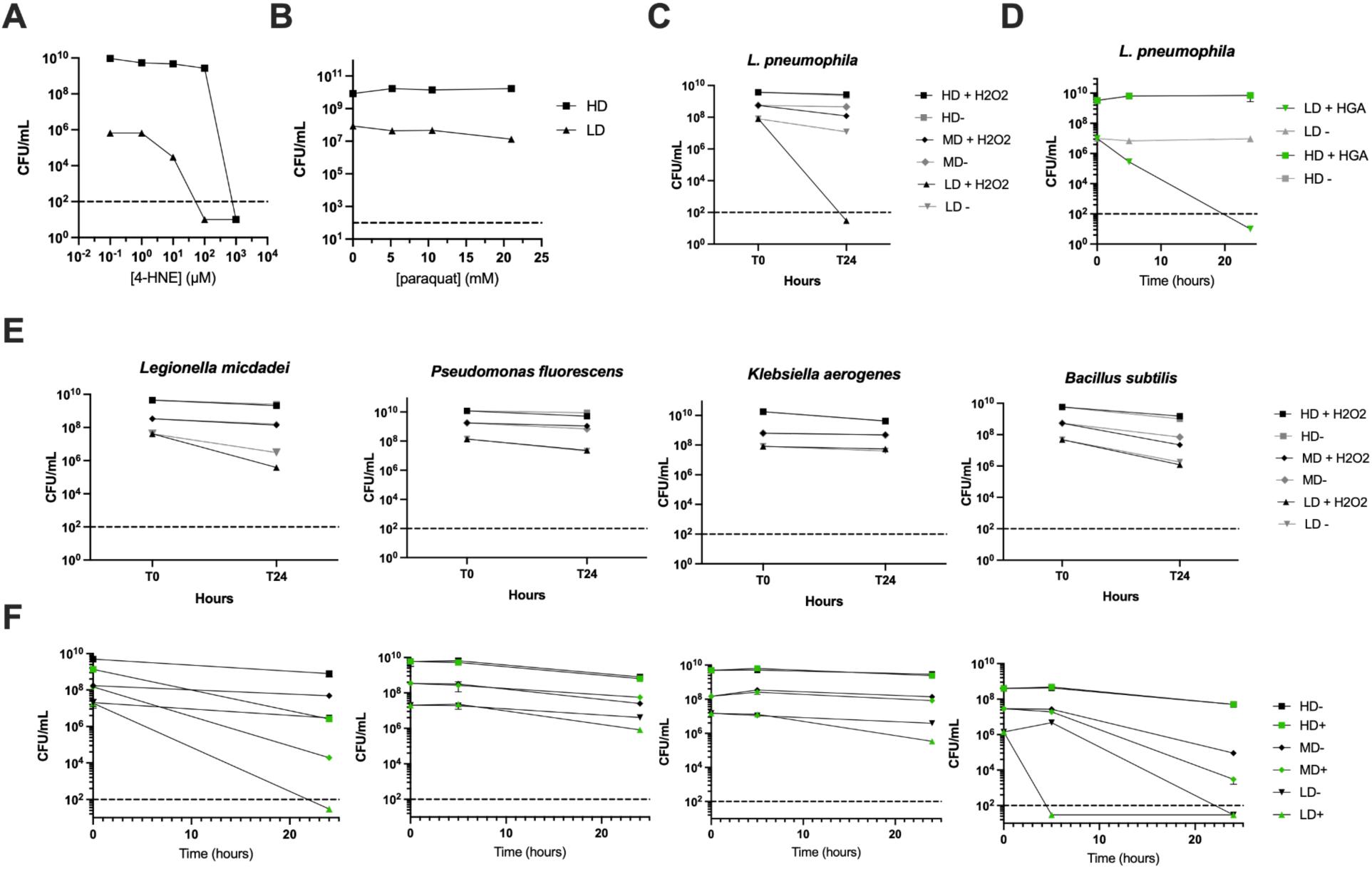
*L. pneumophila’s* density-dependent susceptibility to HGA and H2O2 does not extend to other oxidative stressors nor to other bacterial species. **A)** Similar concentrations of 4-HNE (<10X different) are toxic to high- and low-density *L. pneumophila* **B)** In PBS, *L. pneumophila* is not susceptible to paraquat. **C)** *L. pneumophila* susceptibility to 300 µM H2O2 (compare to E). All species showed minimal sensitivity to H2O2. **D)** *L. pneumophila* susceptibility to 125µM HGA (compare to F) **E-F)** Susceptibility of other bacterial species at high, medium, or low density (HD, MD, LD) to H2O2 **(E)** and HGA**(F)**. *L. micdadei* exhibited some HGA sensitivity at all cell densities. *B. subtilis* had HGA susceptibility at low densities in addition to poor overall viability in PBS.

**Supplemental Figure 3.**
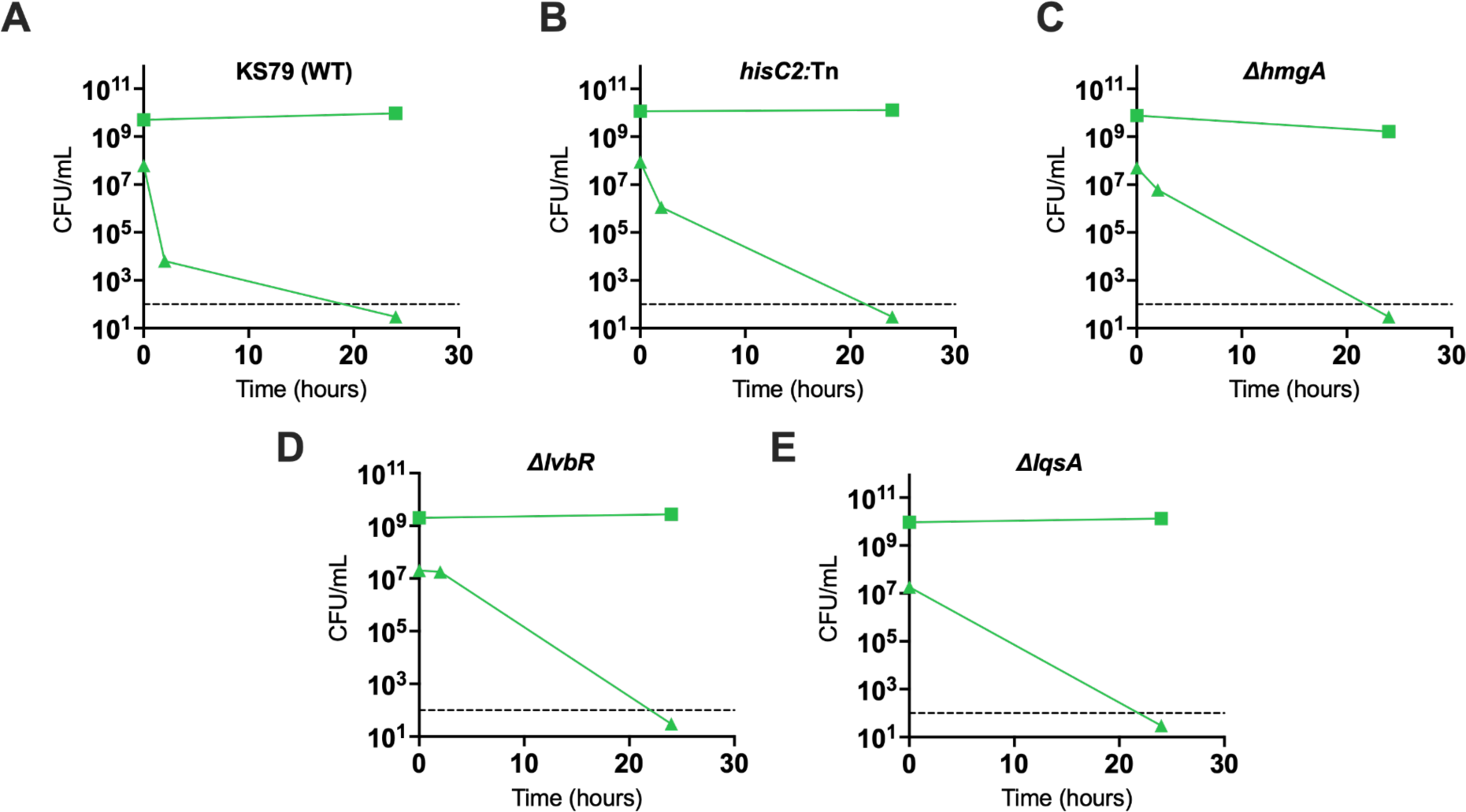
*L. pneumophila’s* density-dependent susceptibility to HGA does not depend on the HGA synthesis or Lqs quorum sensing pathways. **A)** Wild type *L. pneumophila* susceptibilty to 125µM HGA between high-density (square) and low density (triangle) cultures. **B-D)** HGA susceptibilty is unaffected by mutations that eliminate **(B)** or enhance **(C)** HGA production, nor by the deletion of genes encoding the Iqs-regulated transcription factor *IvbR* **(D)** or the Lqs autoinducer synthase *IqsA* **(E).**

**Supplemental Figure 4:**
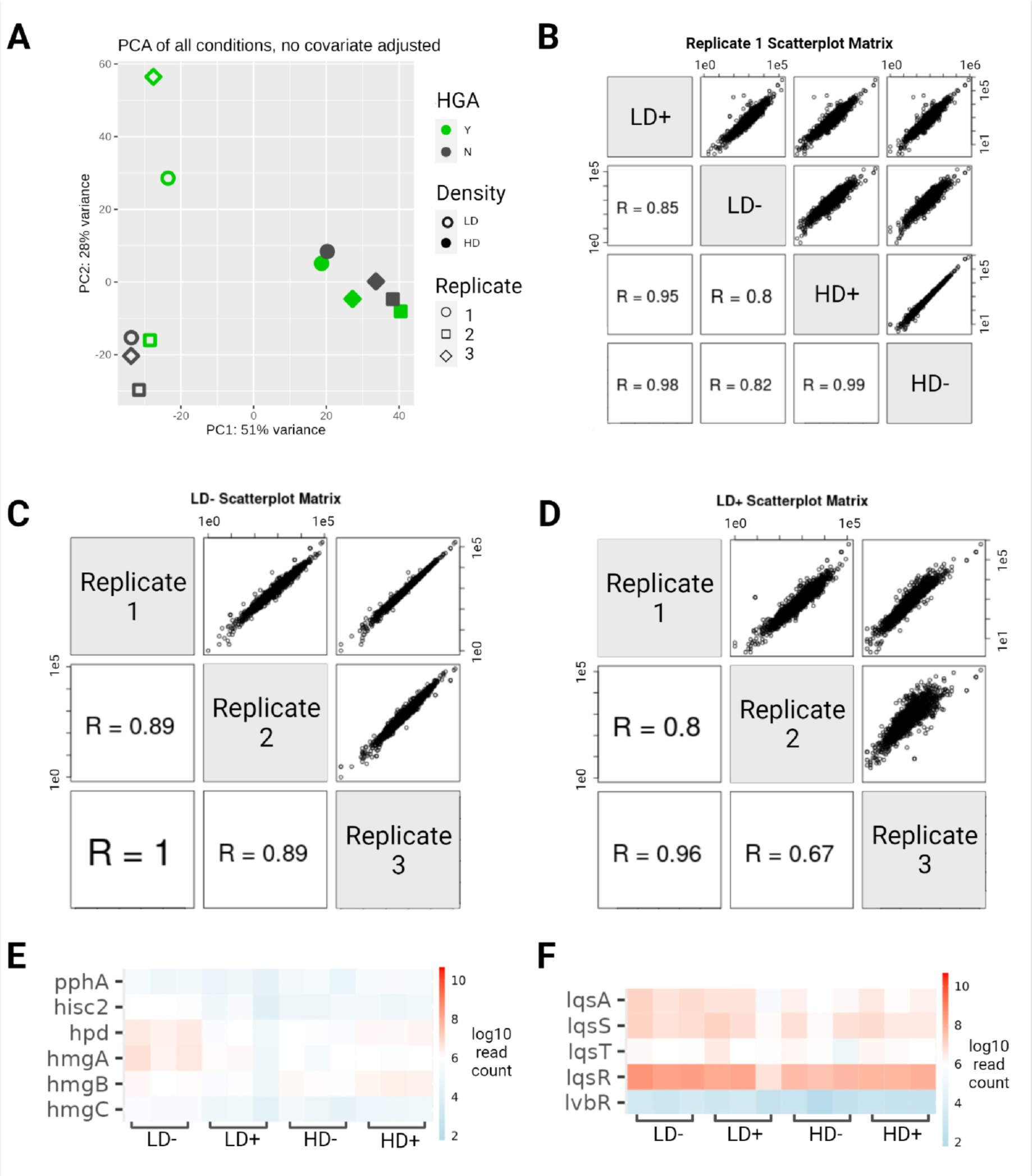
Validation of RNA-seq analyses **A)** Principal component analysis of all RNA-seq samples according to HGA exposure (color), cell density (fill), and replicate (shape). **B-D)** Scatter matrices comparing each gene’s normalized counts within replicate 1 **(B),** between replicates of low density cells without HGA **(C),** and between replicates of low density cells with HGA **(D). E-F)** Heatmaps showing that neither the HGA biosynthesis pathway (E) nor the Legionella quorum sensing(lqs) pathway **(F)** are differentially expressed.

